# Using gene trees with lineage-specific duplicates for phylogenetic inference mitigates the effects of long-branch attraction

**DOI:** 10.1101/2024.12.06.627281

**Authors:** Megan L. Smith, Matthew W. Hahn

## Abstract

Traditionally, the inference of species trees has relied on orthologs, or genes related through speciation events, to the exclusion of paralogs, or genes related through duplication events. This has led to a focus on using only gene families with a single gene-copy per species, as these families are likely to be composed of orthologs. However, recent work has demonstrated that phylogenetic inference using paralogs is both accurate and allows researchers to take advantage of more data. Here, we investigate a case in which using larger gene families actually increases accuracy compared to using single-copy genes alone. Long-branch attraction is a phenomenon in which taxa with long branches may be incorrectly inferred as sister taxa due to homoplasy. The most common solution to long-branch attraction is to increase taxon sampling to break up long branches. Sampling additional taxa is not always feasible, possibly due to extinction or limited access to high-quality DNA. We propose the use of larger gene families with additional gene copies to break up long branches. Using simulations, we demonstrate that using larger gene families mitigates the impacts of long-branch attraction across large regions of parameter space, especially when either maximum parsimony or maximum likelihood with a misspecified substitution model is used for inference. We also analyze data from Chelicerates, with a focus on assessing support for a sister relationship between scorpions and pseudoscorpions. Previous work has suggested that the failure to recover this relationship is due to long-branch attraction between pseudoscorpions and other lineages. Using data from larger gene families increases support for a clade uniting scorpions and pseudoscorpions, although other resolutions of this relationship continue to have higher support. Together, this work highlights the potential utility of larger gene families in phylogenetic inference.

---

A major aim of phylogenetics is to reconstruct evolutionary relationships among species. Advances in sequencing technologies have provided access to thousands of loci, which promises to help resolve recalcitrant nodes in the tree of life (Scornavacca et al., 2020). While phylogenetic methods are powerful, they are sometimes prone to systematic biases, in which incorrect trees are inferred due to unmodelled processes. One of the most well-studied such biases is long-branch attraction (LBA). LBA is a systematic bias in which phylogenetic methods preferentially infer a sister relationship between distantly related taxa with long branches. This occurs because a large number of homoplasious substitutions along long branches leads to high apparent similarity between such taxa (Felsenstein, 1978) (Figure 1A).

**Fig. 1.**
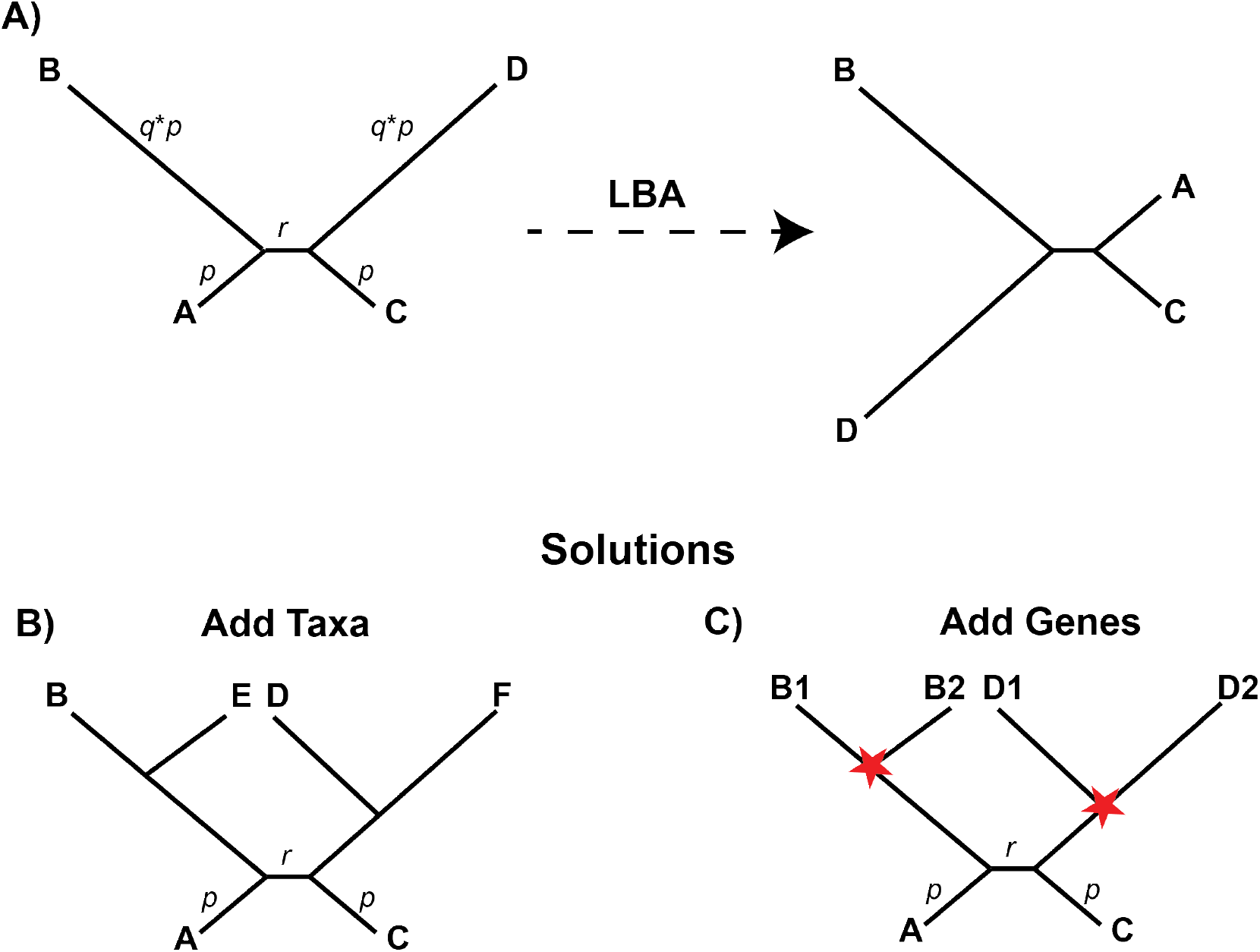
Long-branch attraction (LBA) is a systematic bias that leads phylogenetic methods to incorrectly infer taxa with long branches as sister taxa (A). LBA has often been mitigated by including more taxa to break up long branches (B), but this may not always be possible. Here, we propose using gene duplicates to break up long branches (C). Red stars indicate duplication events. For simulations, we use the tree in A as the species tree.

LBA was first described in the context of parsimony (Felsenstein, 1978; Hendy and Penny, 1989), which is highly susceptible to LBA. Probabilistic approaches—such as maximum likelihood—can somewhat mitigate LBA by accounting for the possibility of homoplasy due to multiple substitutions (Felsenstein, 1978). However, using inadequate substitution models can still leave these methods susceptible to LBA (Gaut and Lewis, 1995; Philippe et al., 2005b; Szánthó et al., 2023). Several solutions have been suggested to mitigate LBA (reviewed in Bergsten 2005), including using more appropriate models of nucleotide substitution (Gaut and Lewis, 1995; Szánthó et al., 2023; Sullivan and Swofford, 1997; Lartillot et al., 2007), excluding quickly evolving sites or genes (Brinkmann et al., 2005; Duchêne et al., 2022; Philippe et al., 2005a; Sanderson et al., 2000), and/or adding taxa to break up long branches (Hendy and Penny, 1989; Graybeal, 1998) (Figure 1B). The latter approach has proven particularly powerful, but sampling additional taxa may be challeging (e.g., due to available resources) or impossible (e.g., due to extinction) in many cases. Here, we explore an alternative approach to breaking up long branches—considering data from larger gene families.

Traditionally, phylogenomics has relied on the use of orthologs (genes related through speciation events) to the exclusion of paralogs (genes related through duplication events). In a typical work flow, researchers begin with a set of coding genes from each sampled taxon. Ortholog clustering approaches (e.g., OrthoMCL; Li et al. 2003) are then used to identify gene families, or groups of closely related genes (sometimes called “orthogroups”). These families include both orthologs and paralogs. The traditional approach is to use only those gene families in which there is a single gene-copy present in each species, as these single copy (SC) genes are hypothesized to be orthologs. Since single-copy orthologs are related only through speciation events, their history should match the species history in the absence of other processes (e.g., incomplete lineage sorting). While this approach has proven powerful in phylogenomics, it results in a massive amount of data loss (Emms and Kelly, 2018). Recent work has explored the potential of using data from larger gene families, either by using tree-based approaches to extract orthologs from these larger gene families (e.g., Yang and Smith 2014; Willson et al. 2022) or by using large gene families directly for inference (e.g., Legried et al. 2021; Zhang et al. 2020, reviewed in Smith and Hahn 2021). Theoretical (Legried et al., 2021; Zhang et al., 2020), simulation (Legried et al., 2021; Zhang et al., 2020; Yan et al., 2022; Parsons and Bansal, 2024), and empirical (Smith et al., 2022) studies have demonstrated that phylogenetic inference using data from larger gene families is highly accurate and allows researchers to use substantially more data.

While previous work has focused on the ability to use more data as the primary reason to analyze larger gene families, here we explore another potential benefit. Just as taxon sampling can be used to break up long branches, gene sampling could also break up long branches (Figure 1C). Specifically, analyzing gene families in which duplication events occur on long branches in the tree should mitigate long-branch attraction. Here, we use simulations to evaluate the impact of including data from larger gene families on inferences using maximum parsimony (MP) and maximum likelihood (ML) in regions of parameter space in which LBA occurs. We explore these effects using both all gene families together and focusing only on gene families with lineage-specific duplications on the long branches of interest. Finally, we analyze data from Chelicerates to evaluate the impacts of including larger gene families on support for a sister relationship between scorpions and pseudoscorpiones, a relationship that has previously been suggested to have been obscured by LBA (Ontano et al., 2021).

## Materials and Methods

### Simulations

We simulated gene families using SimPhy v1.0.2 (Mallo et al., 2015). As a species tree, we used the tree shown in Figure 1A. To vary the strength of long-branch attraction, we varied the length of the internal branch, *r*, and the magnitude of the substitution rate multiplier, *q*. Note that we only modified the substitution rate multiplier, not the external branch lengths themselves. This has practical value, in that it allowed us to input a midpoint-rooted ultrametric tree into SimPhy for conducting simulations. Furthermore, this approach guarantees that only the substitution rate, and not the duplication or loss rates, varies across lineages. Specifically, we explored a grid of values for *r* and *q*. We considered 20 values of each parameter evenly spaced throughout the grid, and we simulated 100 gene trees per parameter combination. In the presence of loss (see below), we may not always have 100 trees per condition, because loss can lead to scenarios in which the four focal species are not all present, and we cannot investigate accuracy in this case. We considered *r* values ranging from 100,000 to 1,000,000 generations. For MP, we considered *q* values ranging from 10 to 30, while for ML, we considered *q* values ranging from 20 to 40 (with a misspecified substitution model; see below) or 40 to 100 with a correctly specified substitution model (because of the greater sensitivity of MP to LBA; Swofford et al. 2001). We set the length of the external branches, *p*, to 250,000 generations. We considered a duplication rate of 1e-6, a nucleotide substitution rate of 1e-7, and a population size of 10,000. We performed two sets of experiments—one with the loss rate set to 0 and a second with the loss rate set equal to the duplication rate. We required a minimum of four leaves in each gene tree when generating data in SimPhy. Values for branch lengths (*r*) and substitution multipliers (*q*) were chosen to ensure that LBA was prevalent across parameter space. Duplication and loss rates were selected to produce reasonable numbers of duplications, single-copy gene families, and gene families with multiple copies in lineages of interest to facilitate relevant comparisons. Finally, we used small population sizes because discordance due to incomplete lineage sorting is not a focus of this study and is not expected to interact with the distribution of duplications under the models implemented in SimPhy.

After simulating gene family trees in SimPhy, we simulated sequence data on each tree using SeqGen v1.3.4 (Rambaut and Grassly, 1997). We used the HKY model of sequence evolution to simulate sequences 2,000 base-pairs in length. We inferred trees from these data using either MP or ML. For MP inference, we used MPBoot v1.1.0 (Hoang et al., 2018) with default settings. For ML inference, we used IQTree v2.3.2 (Minh et al., 2020), and set the nucleotide substitution model to the HKY model used to simulate data.

We also considered a scenario in which ML is expected to be more susceptible to LBA: model misspecification. Parameters were the same as above, except we generated sequence data under a GTR model with gamma-distributed rate variation and a proportion of invariable sites. For gamma-distributed rate heterogeneity, we used an alpha of 5.0 and 4 rate categories. We set the proportion of invariable sites to 0.2. For these data, we inferred gene trees using ML under an overly simplistic HKY model with no rate variation.

To quantify the accuracy of our inferences, we used ASTRAL-IV v1.19.4.5 (Zhang et al., 2018; Zhang and Mirarab, 2022) as implemented in ASTER (available from https://github.com/chaoszhang/ASTER) to calculate quartet concordance factors. These numbers report the fraction of quartets from the inferred gene trees that match the species tree shown in Figure 1A. Since our simulations include only four taxa, the quartet concordance factor is equivalent to the proportion of quartets supporting the species tree topology (averaged across all trees simulated for each condition). In other words, for each gene family, ASTRAL estimates the proportion of quartets that match the species tree topology, returning a value between 0 and 1. We then averaged these values across all replicates within a particular set of conditions. Long-branch attraction will lead to decreased quartet concordance, so we expect concordance factors to be lowest when long-branch attraction has the largest impacts on inference. Notably, gene trees could also differ from the species tree due to incomplete lineage sorting (ILS); however, we expect this to contribute minimally to discordance given the population sizes used here, and the amount of ILS should be constant across data types (e.g., single-copy genes versus all gene families). We also recorded whether each replicate was a single-copy gene, whether the inferred tree included only lineage-specific duplicates (after midpoint rooting), and whether the inferred tree included lineage-specific duplicates on the long branches (after midpoint rooting). Finally, we calculated whether true trees included lineage-specific duplicates on long branches, and, if so, the distance of these duplicates from the furthest leaf.

While these experiments could suggest that lineage-specific duplicates mitigate LBA, several other factors differ across simulation conditions, including the number of quartets and gene families available under each condition. We performed an additional test that held these factors constant. We simulated data as above and focused on gene families with a lineage-specific duplication in at least one long-branch taxon (lineage B or lineage D) and no duplications that were not lineage-specific (i.e., duplications specific to lineage A or lineage C were also allowed). For these gene families, we inferred gene family trees using MP and two different alignments: the full alignment and an alignment with paralogs removed. For the trees inferred from full alignments, we trimmed paralogs from the trees after inference. Then, we calculated quartet concordance from both sets of trees. This analysis should allow us to assess the impacts of lineage-specific duplicates on gene tree estimation error while controlling for other confounding factors.

In addition to these experiments, we performed an experiment in which there was increased discordance due to duplication and loss. In our original simulations, duplications could only occur in the ancestor of a single lineage, in the ancestor of A and B, or in the ancestor of C and D. Because of this, duplications could not generate quartet discordance. On the other hand, if duplications occur in the ancestor of the four focal species, additional discordance may result (Supporting Figure S1), because sampled quartets may include paralogs in multiple lineages. When paralogs are sampled, gene tree quartets may disagree with the species tree even in the absence of gene tree inference error and ILS. Thus, we expect quartet concordance to decrease primarily due to two factors in these simulations: gene tree estimation error and gene duplication and loss. To allow for duplications in the ancestor of the focal species in our simulations, we included an outgroup when simulating data in SimPhy. The outgroup was removed prior to gene tree inference and subsequent analyses. While this can lead to discordance when the full dataset is analyzed using quartet-based approaches, discordance can be avoided by using tree-based decomposition approaches or by sampling only gene families with lineage-specific duplications. For this set of experiments, we varied the ratio between total tree height and the ingroup height, *O* (Supporting Figure S1). We considered *O* values ranging from 1.01 to 5. Because these simulations can lead to much larger trees, we considered duplication and loss rates equal to 5e-7. We set the length of the internal branch *r* equal to 300,000, and allowed *q* to vary as above. All other parameters were set as above. Simulation of sequence data, gene tree inference, and calculations of concordance factors followed the methods described above.

### Chelicerate data

To evaluate whether using paralogs also mitigates LBA in an empirical dataset, we used data from Chelicerates. Previous work has suggested that the lack of support for a sister relationship between pseudoscorpions and scorpions is due to LBA (Ontano et al., 2021), specifically LBA between pseudoscorpions and other long-branch chelicerates. We downloaded 19 whole genomes from NCBI (Sayers et al., 2022), i5k (Thomas et al., 2020), and other sources (Nong et al., 2021; Shingate et al., 2020) (Table S1). For one species, *Tachypleus gigas*, no CDSs were available, so we generated a CDS annotation from the GFF and genome using gffread (Pertea and Pertea, 2020).

Generally, we followed the methodology used to infer gene families from coding sequences used by Thomas et al. (2024). Briefly, we extracted the coding sequence of the longest isoform of each gene using a python script available from https://github.com/gwct/isofilter-gff. We then used FastOrtho with an inflation parameter of 3 (https://github.com/olsonanl/FastOrtho) to cluster genes into gene families. Next, we aligned the sequences of each gene family using mafft v7.526 (Katoh et al., 2009) with a maximum of 5,000 iterations. We then used Gblocks v.0.91b with default parameters to remove poorly aligned positions (Castresana, 2000). Additionally, we used TrimAl to further trim alignments (Capella-Gutiérrez et al., 2009). Finally, we removed all alignments less than 300 base-pairs long.

We inferred gene trees from all Chelicerate alignments using MP and ML. For MP inference, we used MPBoot v1.1.0 (Hoang et al., 2018) with default parameters. For ML inference, we used IQTree v2.3.2 (Minh et al., 2020), selecting the best-fit model automatically using ModelFinder (Kalyaanamoorthy et al., 2017). Finally, to assess support for a sister relationship between the scorpion, *Centruroides sculpturatus*, and the pseudoscorpion, *Cordylochernes scorpioides*, we again used ASTRAL to calculate quartet concordance factors on the tree from Thomas et al. (2024), except that we added *Cordylochernes scorpioides* to the tree as sister to *Centruroides sculpturatus* (Supporting Fig. S2). We calculated concordance factors for six datasets. First, we calculated concordance factors for our trees inferred using MP either including only single-copy genes, all gene families, or only gene families with at least one lineage-specific duplication in pseudoscorpions. Second, we calculated concordance factors for our ML trees for the same three sets of gene families. We calculated confidence intervals by bootstrapping the data. Specifically, we resampled trees from the original dataset with replacement 100 times and calculated concordance factors from each resampled dataset.

## Results

### Using paralogs mitigates long-branch attraction in simulations

We first conducted simulations to demonstrate the effects of long-branch attraction when using single-copy genes. As expected, the number of single-copy genes is highest when the internal branch *r* is short, as there is less time for gene duplication events (Supporting Fig. S3). Additionally, larger values of *r* lead to relatively fewer lineage-specific duplicates (LSDs), as there is more time for duplications in the ancestors of (A,B) and (C,D) (Supporting Fig. S3). The overall number of observed duplications also increases with increasing *r*, as expected. Across all of the MP simulations, the average number of observed duplications (i.e., those with at least one copy surviving to the present) was 2.18 without loss and 2.21 with loss. The average number of gene copies (i.e., leaves) per gene family was 6.8 without loss and 5.8 with loss. As expected, the accuracy of inferred trees decreases as the length of the internal branch (*r*) decreases, and as the branch length multiplier (*q*) increases (Fig. 2). This pattern is observed under all conditions, but is much more pronounced under MP (Fig. 2A) and under ML with a misspecified substitution model (Fig. 2G) than under ML with a correctly specified substitution model (Fig. 2D). (Note the difference in the *q* -values used on the y-axes across rows.)

**Fig. 2.**
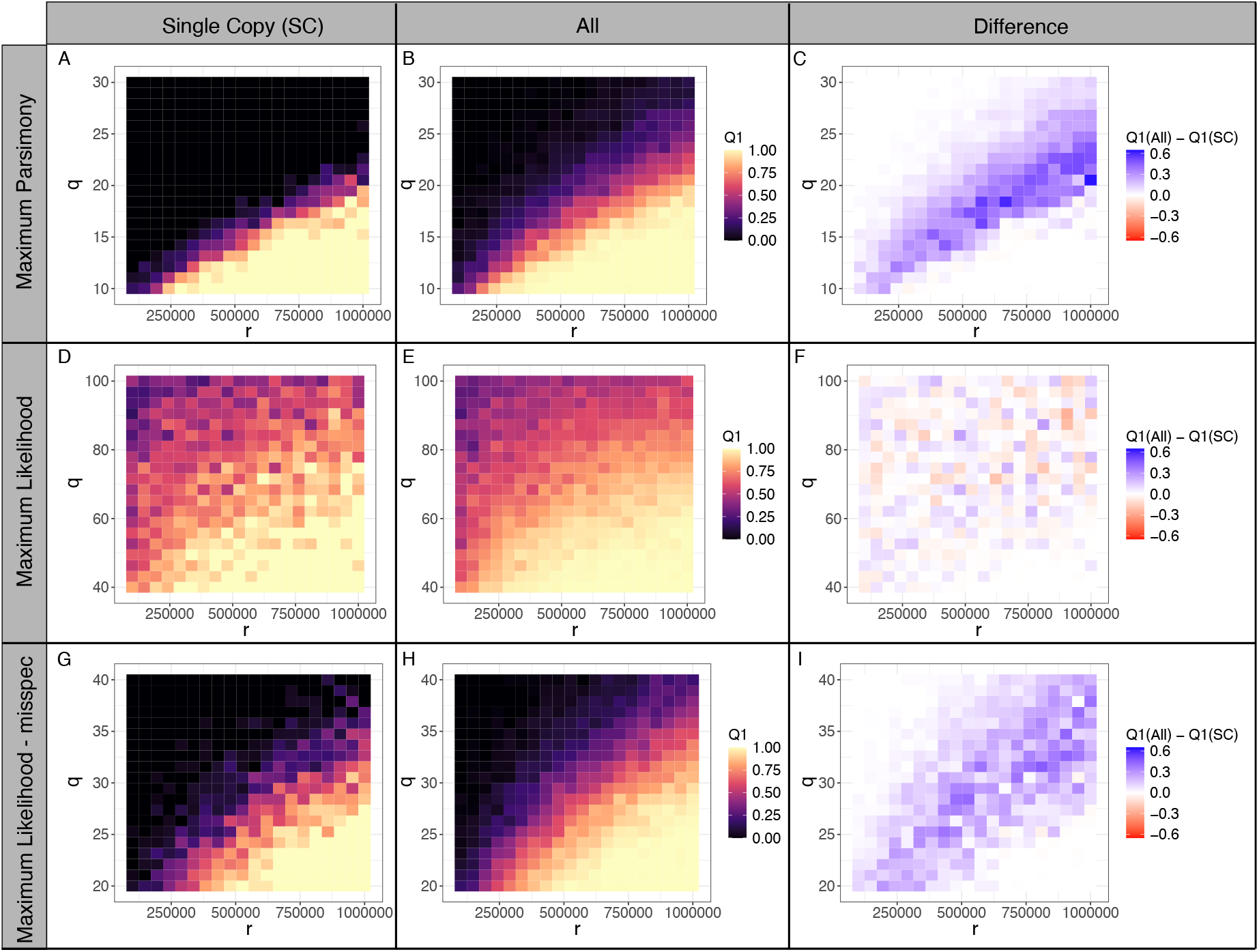
Quartet concordance with the species tree (Q1) across values of the long branch multiplier, *q*, and the internal branch length, *r*. A) Maximum parsimony (MP) with single copy (SC) genes. B) MP with all gene families. C) Difference in Q1 between All and SC genes using MP. D) ML with SC genes under the correct model of sequence evolution. E) ML with all gene families under the correct model of sequence evolution. F) Difference in Q1 between All and SC genes using ML under the correct model of sequence evolution. G) ML with SC genes under a misspecified model of sequence evolution. H) ML with all gene families under a misspecified model of sequence evolution. I) Difference in Q1 between All and SC genes using ML under a misspecified model of sequence evolution. Parameter values used in simulations: *λ*=1e-6; *µ*=0; *p*=250,000; substitution rate = 1e-7; *N*_*e*_ = 10000; *L* = 2000

Next, we inferred trees using data from all gene families, which includes duplicates that can break up the long branches. Using all gene families resulted in increased accuracy for both MP (Fig. 2B,C) and ML with a misspecified substitution model (Fig. 2H,I). In general, the accuracy stayed high in regions of parameter space in which accuracy was already high when using single-copy genes, but using all gene families extended the parameter space across which accurate trees could be inferred. Results were less clear for ML with a correctly specified model, as changes in concordance appeared random across much of parameter space (Fig. 2E,F). The above results are all for simulations with no gene loss, but the same trends held when the loss rate, *µ*, was set equal to the duplication rate, *λ* (Supporting Fig. S4).

Duplicates that occur very near the present may be less effective at breaking up long branches compared to duplicates that occur at intermediate ages. To further explore the role of lineage-specific duplicates in mitigating LBA, we calculated the maximum distance between a lineage-specific duplication node along the lineages leading to species B or D (the long branches) to the furthest leaf node on simulated gene trees. We found that the age of the oldest LSD was larger for gene families for which inferred trees were concordant with the species tree (Fig. 3). Analyzing only gene families with LSDs with and without trimming paralogs from alignments prior to inference further supports that LSDs mitigate LBA. When paralogs are trimmed from alignments, results from gene families with LSDs are similar to those from only SC genes (Supporting Fig. S5A). However, when gene family trees are inferred from alignments that include LSDs (and those LSDs are trimmed prior to calculating quartet concordance), there is a substantial increase in gene tree accuracy (Supporting Fig. S5B,C).

**Fig. 3.**
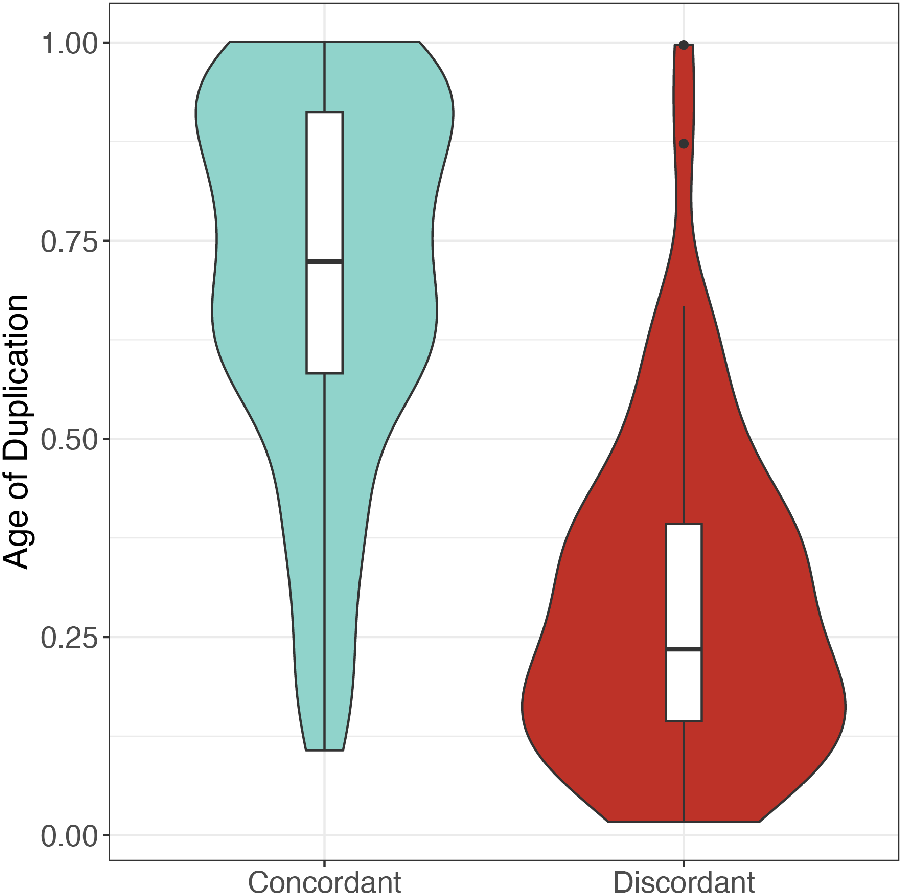
The age (calculated as the distance to the furthest leaf divided by the total branch length) of the oldest duplicate on branches leading to the long branch taxa B and D. Only gene families with lineage-specific duplicates on the long branches were included. LSDs were identified and ages were calculated based on true gene trees, while concordance was assessed based on inferred gene trees. Parameter values used in simulations: *λ*=1e-6; *µ*=0; *p*=250,000; substitution rate = 1e-7; *N*_*e*_ = 10000; *L* = 2000; *q* = 20; *r* = 750, 000.

Finally, we explored the impact of adding duplications in the ancestor of the focal taxa, which can increase discordance due to gene duplication and loss (i.e., pseudoorthologs; Smith and Hahn 2021). As expected, these duplications have no impact when analyzing single-copy genes, as paralogs are unlikely to be included. Specifically, the amount of concordance observed in Fig. 4A is equivalent to that seen in 2A with a fixed *r* of 750,000. However, when analyzing all gene families, additional opportunities for discordance due to duplication and loss generally lead to lower quartet concordance. The larger the value of *O* —i.e. the longer the branch subtending the ingroup—the higher the probability of duplications on that branch. At small to medium values of *q*, increasing *O* leads to more discordance (Fig. 4B,E,H, Supporting Fig. S6A,C,E). The situation is more complicated when *q* is large. Under MP and ML with a misspecified substitution model, when *q* is large, the systematic biases due to long-branch attraction lead to concordance factors near zero in the absence of paralogs (Fig. 4A,G). In this case, adding noise due to paralogs actually increases the proportion of quartets that agree with the species tree. Thus, the added noise due to duplication and loss increases concordance relative to that seen in the case of extreme systematic bias (Fig. 4B,H; Supporting Fig. S6A,E).

**Fig. 4.**
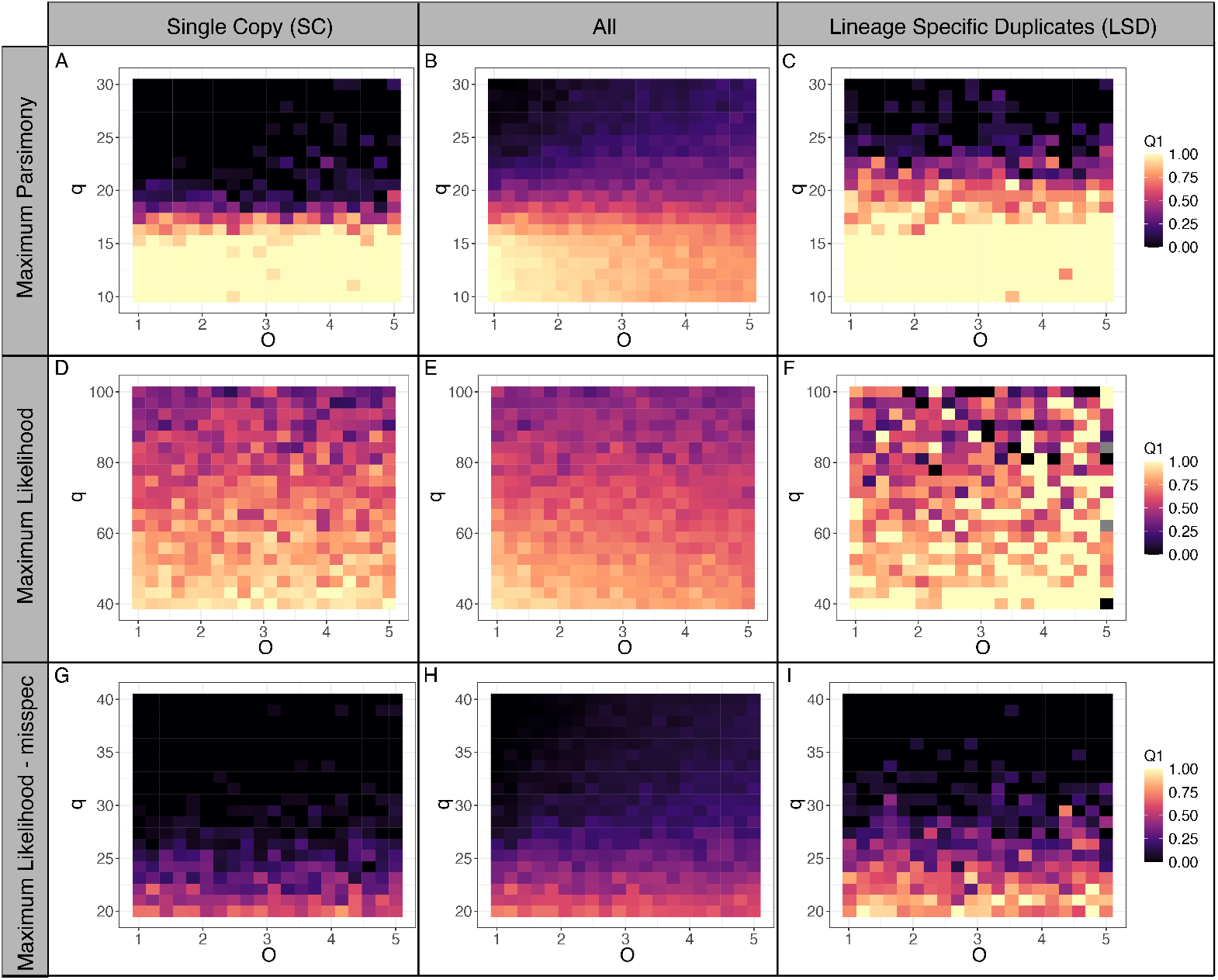
Quartet concordance across values of the long branch multiplier, *q*, and the ratio between total tree height and the ingroup height, *O*. A) Maximum parsimony (MP) with single copy (SC) genes. B) MP with all gene families. C) MP with gene families that include a lineage-specific duplicate in the long-branch taxa and no duplications that are not lineage-specific. D) ML with SC genes under the correct model of sequence evolution. E) ML with all gene families under the correct model of sequence evolution. F) ML with gene families that include a lineage-specific duplicate in the long-branch taxa and no duplications that are not lineage-specific under the correct model of sequence evolution. G) ML with SC genes under a misspecified model of sequence evolution. H) ML with all gene families under a misspecified model of sequence evolution. I) ML with gene families that include a lineage-specific duplicate in the long-branch taxa and no duplications that are not lineage-specific under a misspecified model of sequence evolution. LSDs were identified based on inferred gene trees. Parameter values used in simulations: *λ*=5e-7; *µ*=5e-7; *p*=250,000; *r* = 750,000 for MP and r = 300,000 for ML; substitution rate = 1e-7; *N*_*e*_ = 10000; *L* = 2000

To avoid issues with duplication and loss complicating relationships, researchers need not rely only on single-copy genes. Instead, researchers can filter their data to include only lineage-specific duplicates. Furthermore, if the goal is to mitigate LBA, then using only those gene families with gene duplications on the long branches should be particularly effective (Fig. S5). Notably, no knowledge of true gene family trees is needed to do this: researchers can use simple filtering approaches to include only gene families with two gene copies in the taxa of interest. When we calculated accuracy using only those gene families with relevant lineage-specific duplicates, concordance was again increased relative to the single-copy genes using both MP and ML with a misspecified substitution model (Fig. 4C,I; Supporting Figure S6B,F). As above, there was no clear pattern when ML was used with the correct substitution model (Fig. 4F; Supporting Figure S6D).

### Including data from larger gene families increases support for a sister relationship between pseudoscorpions and scorpions

LBA has been proposed to decrease support for a sister relationship between scorpions and pseudoscorpions (Ontano et al., 2021), so we expect that support for this relationship will be higher under ML than under MP (since MP is more susceptible to LBA). Our results support this prediction, with more trees supporting their sister relationship when inferred using ML, though the confidence intervals are overlapping (“SC” row in Table 1). Furthermore, if using larger gene families helps to mitigate LBA, then we expect that support for this relationships will be higher when all gene families are analyzed compared to when only single-copy genes are analyzed. Again, our results support this prediction, with higher support for sister relationships between scorpions and pseudoscorpions when all gene families are used (“All” row in Table 1); however, confidence intervals are again overlapping with estimates from single-copy datasets.

**Table 1.** Quartet concordance for a sister relationship between pseudoscorpions and scorpions across maximum parsimony (MP) and maximum likelihood (ML) datasets with only single copy genes (SC), all gene families (All), and only gene families with a lineage-specific duplication on the branch leading to pseudoscorpions (LSD).

| Data | MP (95% CI) | ML (95% CI) | # gene families |
| --- | --- | --- | --- |
| SC | 0.146 (0.119, 0.174) | 0.187 (0.157, 0.216) | 2787 |
| All | 0.157 (0.139, 0.177) | 0.198 (0.178, 0.215) | 4348 |
| LSD | 0.154 (0.120, 0.197) | 0.205 (0.164, 0.251) | 187 (MP), 183 (ML) |

Finally, we expect support for this sister relationship to be highest when we only analyze lineage-specific duplicates, as this should allow us to break up long branches without adding noise due to gene duplication and loss. While our results support this prediction when using ML (“LSD” row in Table 1), the same was not true under MP, and confidence intervals always overlapped with those found when analyzing all gene families. Notably, relatively few gene families with lineage specific duplicates were present in this dataset (Table 1). Despite this promising trend, regardless of how we analyze these data, relationships other than a sister relationship between scorpions and pseudoscorpions are supported by the data (Supporting Table S2).

## Discussion

Traditionally, phylogenetic inference has relied on orthologs, primarily by restricting analyses to single-copy genes. Recent work has demonstrated that using data from larger gene families allows researchers to use more data without sacrificing accuracy (Smith et al., 2022; Yan et al., 2022; Legried et al., 2021; Zhang et al., 2020; Parsons and Bansal, 2024). While this work has generally focused on the advantages of using larger gene families when single-copy genes are limited in number, here we focus on a case in which individual gene trees inferred from larger gene families have increased accuracy.

LBA appears to be common across the tree of life (e.g., Ontano et al. (2021); Cai et al. (2024); Szánthó et al. (2023)). While several solutions have been proposed, no single solution is without its challenges. Adding more taxa is an effective solution (Hendy and Penny, 1989; Graybeal, 1998), but is not always possible due to extinction or a lack of available samples. Using more complex substitution models has also proven powerful in many contexts, particularly when taking into account heterogeneity in the substitution process (Lartillot et al., 2007; Szánthó et al., 2023). However, estimating parameters of complex models that account for rate heterogeneity (e.g., CAT models; Lartillot and Philippe 2004) is challenging and computationally expensive, and may not always result in more accurate inferences (Whelan and Halanych, 2017). Thus, novel approaches to mitigating LBA are still needed.

Many recent approaches to accurate phylogenetic inference combine data from multiple gene trees to infer a species tree (e.g., ASTRAL, Zhang and Mirarab 2022; ASTRID, Vachaspati and Warnow 2015). These methods are explicitly avoiding biases associated with concatenating gene trees (Kubatko and Degnan, 2007), but may themselves suffer from issues inherent in inferring gene trees from short alignments (Bryant and Hahn, 2020). If gene trees are not inferred accurately, then the species tree may also not be inferred accurately (Molloy and Warnow, 2018), especially for branches deep in time (Rosenzweig et al., 2022). In fact, both types of methods—those that use gene trees and those that use concatenated ML—can fail due to systematic biases, like LBA (Roch et al., 2019). Using data from larger gene families offers the potential for mitigating LBA in individual gene trees with minimal costs. In particular, using gene families with only lineage-specific duplicates makes it possible to break up long branches without adding any additional noise due to gene duplication and loss. Our simulation studies support the idea that using such gene families should increase accuracy without increasing noise in the case of MP and ML with a misspecified substitution model (Fig. 4; Supporting Figure S5), hopefully also making the inference of species trees from such gene trees more accurate.

Of course, some empirical datasets may have few lineage-specific duplicates, as seen in the Chelicerate dataset analyzed here. However, other datasets will have a plethora of such gene families (e.g., primates; Smith et al. 2022). We anticipate that this approach will be more useful in cases in which more such gene families are available. If duplication rates, as well as substitution rates, are elevated in long-branch taxa, we might expect more relevant gene families to be present. Regardless, we expect that this approach will be helpful in some empirical cases plagued by LBA.

Notably, improvements are only clear in cases in which LBA is expected to be a major problem in inference—either using MP or using ML with an overly simplistic substitution model. While MP has become less popular in recent years, ML is a major workhorse for phylogenetic inference. In the relatively simple case considered here, ignoring gamma-distributed rate heterogeneity and a proportion of invariable sites, as well as misspecifying the substitution model, led to large amounts of LBA (Fig. 2G). While we could have analyzed these data under a more appropriate model, using (or estimating) appropriately complex substitution models is not always possible with empirical datasets. Our results suggest that, when ML is susceptible to LBA due to a misspecified substitution model, using data from larger gene families offers a potential path toward more accurate inference.

The simulation studies conducted here focus on a relatively simple model of gene duplication and loss. Notably, we do not consider variation in gene duplication and loss rates across lineages or gene families. We expect that variation in rates across gene families would primarily impact which gene families were most likely to have helpful lineage-specific duplicates. On the other hand, variation across lineages could have mixed impacts. Higher rates of duplication on long-branch lineages could lead to more lineage-specific duplicates, thus further ameliorating the impacts of LBA. It is less clear to us how varying loss rates might impact this approach. Certain patterns of differential loss could lead to an increase in the proportion of pseudoorthologs, which are rare under simpler models (Smith and Hahn, 2022). It is possible to imagine a complex model in which differential patterns of loss could mislead inference, but to do so requires a scenario that may or may not be biologically realistic. Future work should evaluate the impact of more complex models of duplication and loss on phylogenetic inference using data from larger gene families.

Our empirical analysis of chelicerates was less decisive. While support for pseudoscorpions and scorpions as sister lineages increased when using all gene families or lineage-specific duplicates, the increase was not large. This may be due in part to the relatively small size of our datasets, particularly the dataset including only lineage-specific duplicates: only 183-187 genes (depending on the method used) showed lineage-specific duplicates in pseudoscorpions (Table 1). In addition, though LBA was proposed to be the cause of phylogenetic uncertainty in the chelicerates (Ontano et al., 2021), this is by no means certain. Applications to more systems in which LBA is thought to act (e.g. Bergsten 2005) would offer additional tests of our method.

Our results suggest a potential path forward to mitigating long-branch attraction using genomic data. Future research should explore the impacts of other methods for inferring trees from larger gene families, including methods that do not simply use all orthologs and paralogs together. Methods that either account for gene duplication and loss explicitly (e.g., ASTRAL-Pro; Zhang et al. 2020) or decompose larger trees into more manageable sub-trees (Yang and Smith, 2014; Smith et al., 2022) could further remove noise even while taking advantage of duplicated genes. We anticipate that applying such approaches would provide more ways to mitigating LBA using data from larger gene families.

## Acknowledgements

This work was supported by a National Science Foundation grant to M.W.H. (DBI-2146866).

## Data Availability

All scripts and input files used in this study are available at https://github.com/SmithLabBio/paralogs_lba and as a stable release through Zenodo (https://doi.org/10.5281/zenodo.16929118)

## Notes

### Competing Interest Statement

The authors have declared no competing interest.

### Summary of Updates

New analyses were added to highlight an additional scenario in which paralogs can mitigate LBA: when maximum likelihood is used with a misspecified substitution model. We also added an analysis where we analyzed gene families with lineage-specific duplicates either removing or retaining paralogs.

