## Supporting Information for "Using gene trees with lineage-specific duplicates for phylogenetic inference mitigates the effects of long-branch attraction"

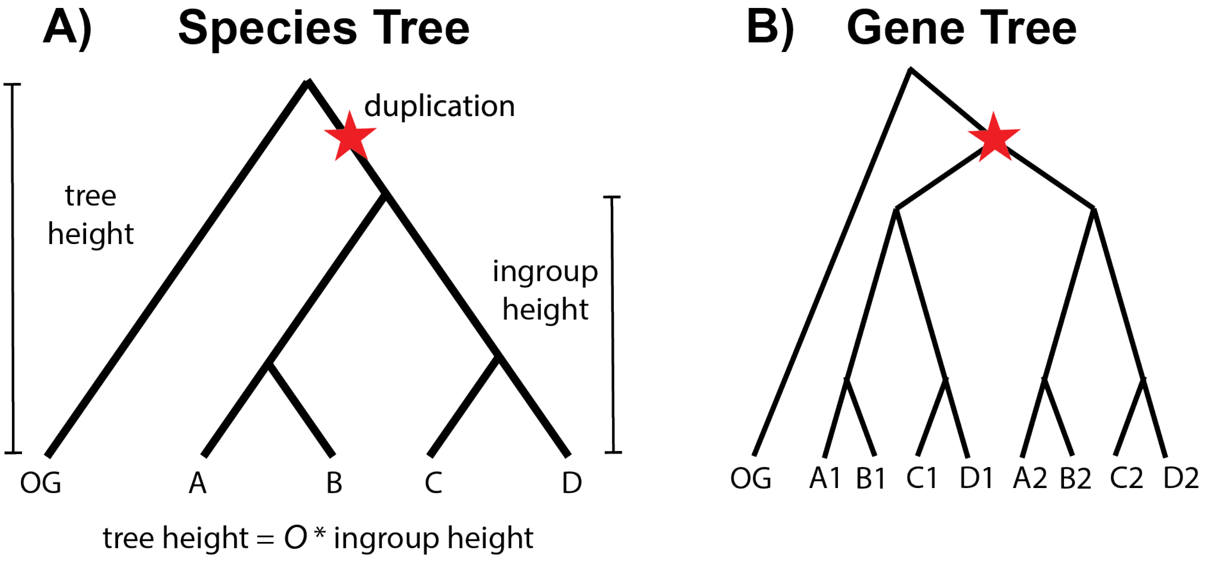


**C) Sampled Quartets**

**Orthologs:**
((A1,B1),(C1,D1));
((A2,B2),(C2,D2));

**Paralogs (Match Species Tree):**
((A1,B1),(C1,D2));
((A2,B2),(C2,D1));
((A1,B1),(C2,D1));
((A2,B2),(C1,D2));
((A1,B2),(C1,D1));
((A2,B1),(C2,D2));
((A2,B1),(C1,D1));
((A1,B2),(C2,D2));
((A1,B1),(C2,D2));
((A2,B2),(C1,D1));

**Paralogs (Alternate 1):**
(A1,C1),(B2,D2));
(A2,C2),(B1,D1));

**Paralogs (Alternate 2):**
(B1,C1),(A2,D2));
(B2,C2),(A1,D1));

Fig. S1. When data are simulated on a species tree including an outgroup (A), duplications can occur in the ancestor of the focal taxa (B). This leads to discordance, as evidenced by looking at the quartets that can be sampled in this case (C). When sampling quartets for a set of species (A,B,C,D), we sample a single copy from each species. In some cases, we sample four orthologs. However, we often sample paralogs. These paralogs can either have topologies that match the species tree, or topologies that disagree with the species tree (Alternate 1 or Alternate 2). Red stars indicate gene duplication events.

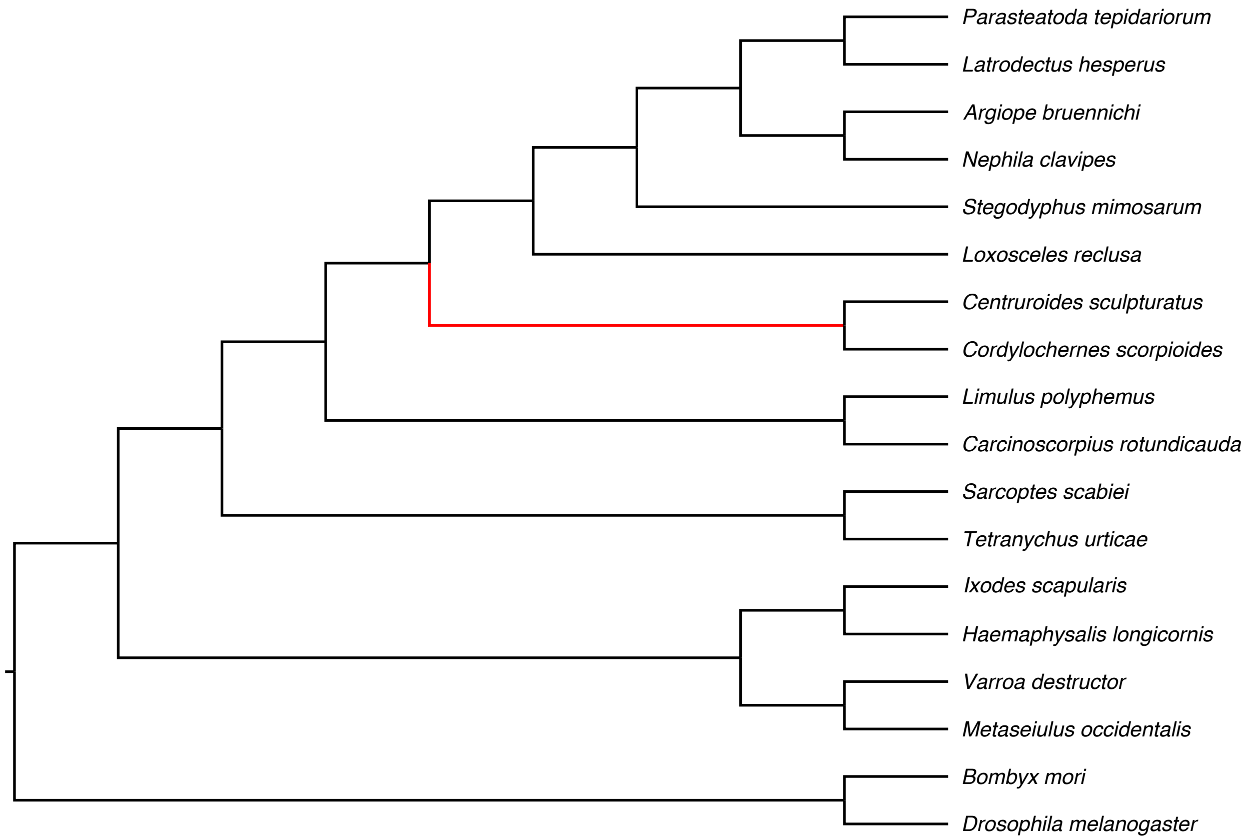


Fig. S2. Tree used for assessing support for Chelicerate relationships.

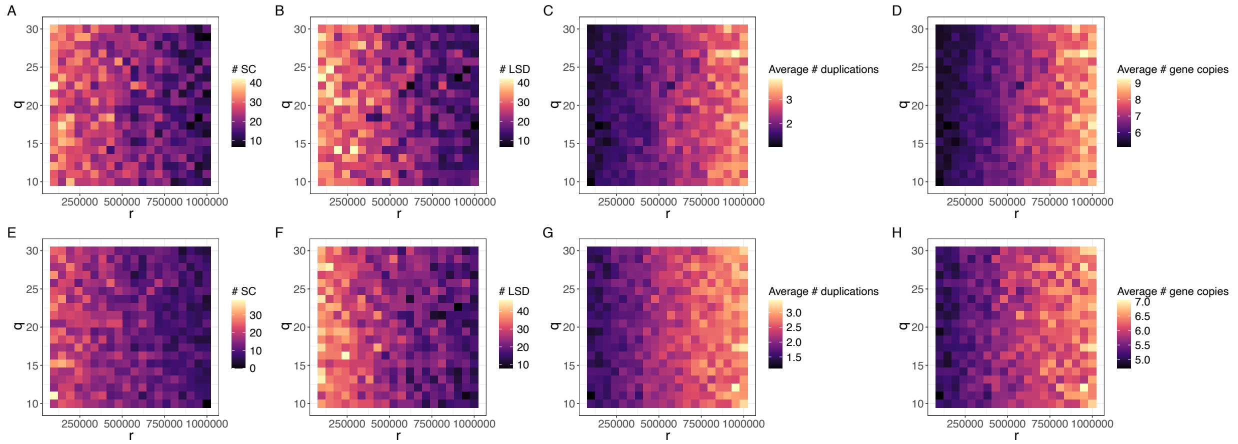


Fig. S3. Data set characteristics under the Maximum Parsimony simulations with and without loss. A) Number of single copy (SC) gene families per grid cell without loss. B) Number of gene families with lineage-specific duplicates on lineages B and D and no duplications that aren't lineage specific without loss. C) Average number of observed duplications without loss. D) Average number of gene copies per gene family without loss. E) Number of single copy (SC) gene families per grid cell with loss. F) Number of gene families with lineage-specific duplicates on lineages B and D and no duplications that aren't lineage specific with loss. G) Average number of observed duplications with loss. H) Average number of gene copies per gene family with loss. To identify lineage-specific duplicates, we used true tree topologies. Parameter values used in simulations: $\lambda$=1e-6; $\mu$=0 or 1e-6; $p$=250,000; substitution rate = 1e-7; $N_e$ = 10000; $L$ = 2000

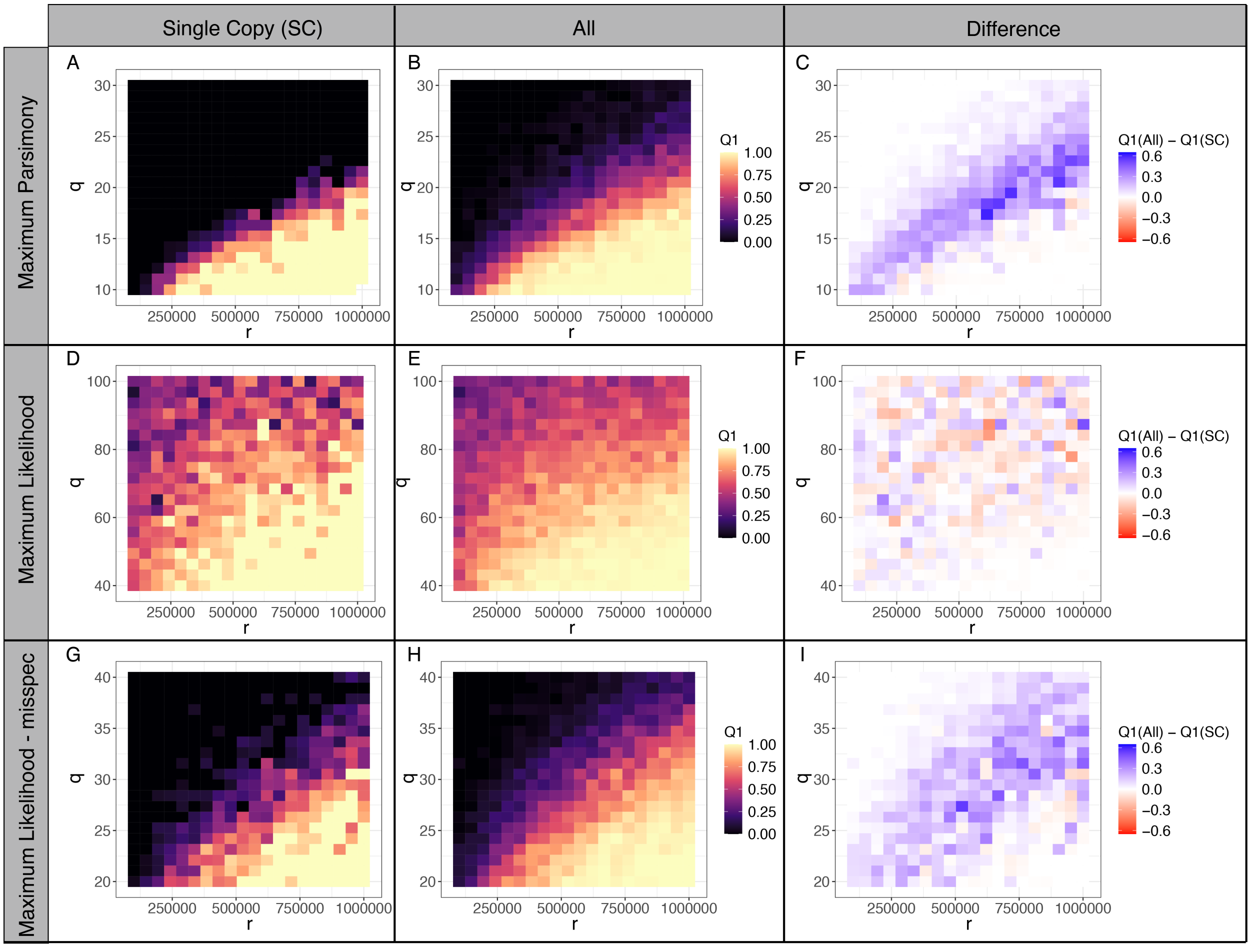


Fig. S4. Quartet concordance across values of the long branch multiplier, $q$, and the internal branch length, $r$. A) Maximum parsimony (MP) with single copy (SC) genes. B) MP with all gene families. C) Difference in Q1 between All and SC genes using MP. D) ML with SC genes under the correct model of sequence evolution. E) ML with all gene families under the correct model of sequence evolution. F) Difference in Q1 between All and SC genes using ML under the correct model of sequence evolution. G) ML with SC genes under a misspecified model of sequence evolution. H) ML with all gene families under a misspecified model of sequence evolution. I) Difference in Q1 between All and SC genes using ML under a misspecified model of sequence evolution. Parameter values used in simulations: $\lambda$=1e-6; $\mu$=1e-6; $p$=250,000; substitution rate = 1e-7; $N_e$ = 10000; $L$ = 2000

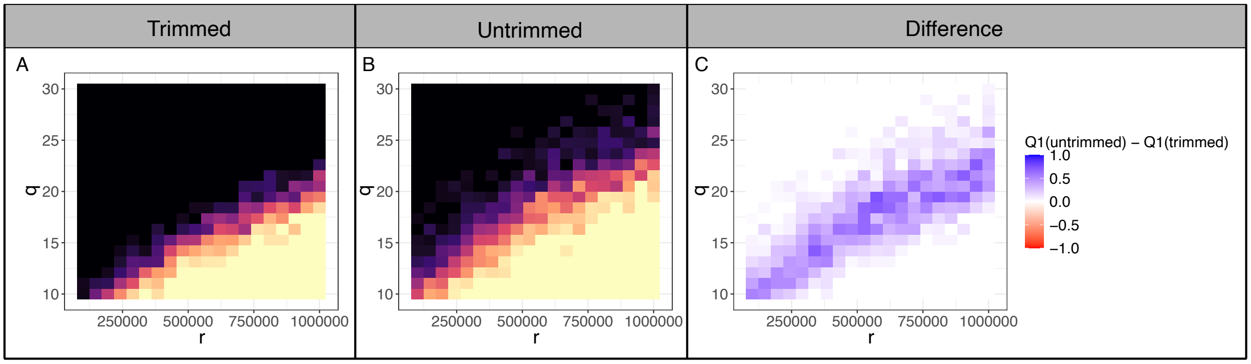


Fig. S5. Quartet concordance across values of the long branch multiplier, $q$, and the internal branch length, $r$ when analyzing gene families with lineage-specific duplicates (LSDs). A) Results on gene trees estimated from alignments in which paralogs were trimmed prior to gene tree inference. B) Results on gene trees estimated from alignments including paralogs, with paralogs trimmed only after gene tree inference. C) Difference in Q1 between the untrimmed and trimmed datasets. Parameter values used in simulations: $\lambda$=1e-6; $\mu$=0; $p$=250,000; substitution rate = 1e-7; $N_e$ = 10000; $L$ = 2000

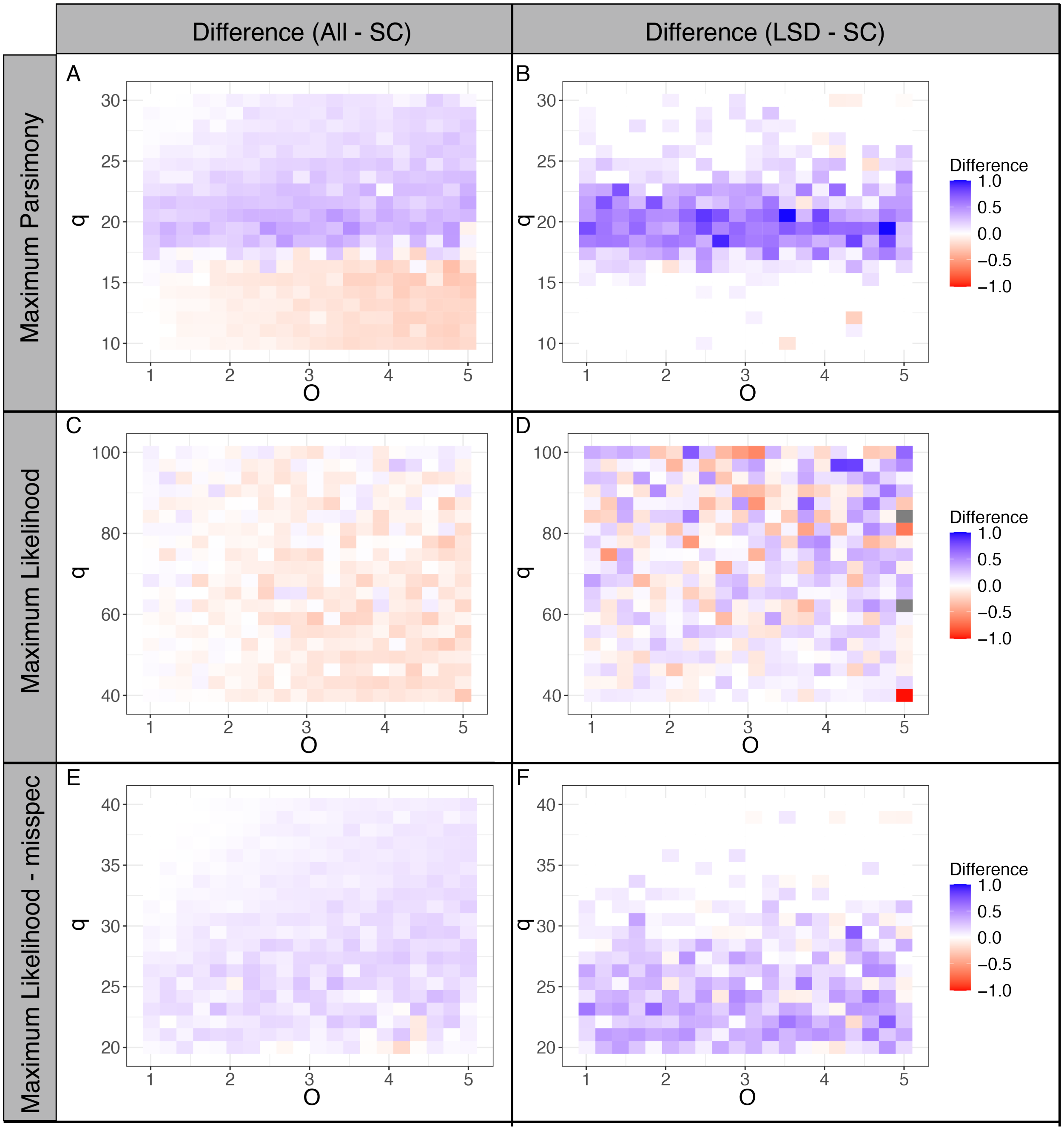


Fig. S6. Difference in quartet concordance across values of the long branch multiplier, $q$, and the ratio between total tree height and the ingroup height, $O$. A) Difference in Q1 between All and single copy (SC) genes using MP. B) Difference in Q1 between lineage-specific duplicates (LSD) and SC genes using MP. C) Difference in Q1 between All and SC genes using Maximum Likelihood (ML) under the correct model of sequence evolution. D) Difference in Q1 between LSD and SC genes using Maximum Likelihood (ML) under the correct model of sequence evolution. E) Difference in Q1 between All and SC genes using Maximum Likelihood (ML) under a misspecified model of sequence evolution. F) Difference in Q1 between LSD and SC genes using Maximum Likelihood (ML) under a misspecified model of sequence evolution. LSDs were identified based on inferred gene trees. Parameter values used in simulations: $\lambda$=5e-7; $\mu$=5e-7; $p$=250,000; $r$ = 750,000 for MP and r = 300,000 for ML; substitution rate = 1e-7; $N_e$ = 10000; $L$ = 2000

| Species name | NCBI Accession | Source | Link (non-NCBI) | Notes |
|---|---|---|---|---|
| Argiope bruennichi | GCA_015342795.1 | NCBI | | |
| Bombyx mori | GCA_000151625.1 | NCBI | | |
| Carcinoscorpius rotundicauda | | FigShare | https://doi.org/10.6084/m9.figshare.13172414.v2 | |
| Cordylochernes scorpioides | GCA_030710605.1 | NCBI | | |
| Centruroides sculpturatus | GCA_000671375.2 | NCBI | | |
| Drosophila melanogaster | GCA_000001215.4 | NCBI | | |
| Haemaphysalis longicornis | GCA_013339765.2 | NCBI | | |
| Ixodes scapularis | GCA_000208615.1 | NCBI | | |
| Latrodectus hesperus | | i5k | https://i5k.nal.usda.gov/content/data-downloads | Current Genome Assembly Official or Primary Gene Set BCM_version_0.5.3 consensus_gene_set |
| Limulus polyphemus | GCA_000517525.1 | NCBI | | |
| Loxosceles reclusa | | i5k | https://i5k.nal.usda.gov/content/data-downloads | Current Genome Assembly Official or Primary Gene Set BCM_version_0.5.3 consensus_gene_set |
| Metaseiulus occidentalis | GCA_000255335.2 | NCBI | | |
| Nephila clavipes | GCA_002102615.1 | NCBI | | |
| Parasteatoda tepidariorum | GCA_000365465.3 | NCBI | | |
| Stegodyphus mimosarum | GCA_000611955.2 | NCBI | | |
| Sarcoptes scabiei | GCA_000828355.1 | NCBI | | |
| Tachypleus gigas | | Dryad | https://doi.org/10.5061/dryad.2jm63xsmc | |
| Tetranychus urticae | GCA_000239435.1 | NCBI | | |
| Varroa destructor | GCA_002443255.1 | NCBI | | |

Table S1. Whole genomes used for the Chelicerate analyses.

| Data | MP: Q1, Q2, Q3 | ML: Q1, Q2, Q3 | # gene families |
|---|---|---|---|
| SC | 0.146, 0.156, 0.699 | 0.187, 0.172, 0.641 | 2787 |
| All | 0.157, 0.184, 0.659 | 0.198, 0.193, 0.609 | 4348 |
| LSD | 0.154, 0.195, 0.651 | 0.205, 0.211, 0.583 | 187 (MP), 183 (ML) |

Table S2. Quartet concordance for all three alternate resolutions across maximum parsimony (MP) and maximum likelihood (ML) datasets with only single copy genes (SC), all gene families (All), and only gene families with a lineage-specific duplication on the branch leading to pseudoscorpions (LSD).
